# Interpretation of the morphological adaptations associated with viviparity in the Tsetse fly (*Glossina morsitans*) by three dimensional analysis

**DOI:** 10.1101/2020.06.12.147587

**Authors:** GM Attardo, N Tam, DY Parkinson, LK Mack, XJ Zahnle, J Arguellez, P Takáč, AR Malacrida

## Abstract

Tsetse flies (genus *Glossina*), the sole vectors of African trypanosomiasis, are distinct from other disease vectors, and most other insects, due to dramatic morphological and physiological adaptations required to support their unique life histories. These evolutionary adaptations are driven by demands associated with their strict dietary and reproductive requirements. Tsetse reproduce by obligate viviparity which entails obligate intrauterine larval development and provisioning of nutrients for the developing larvae. Viviparous reproduction reduces reproductive capacity/rate which also drives increased inter- and intra-sexual competition. Here, we use phase contrast microcomputed tomography (pcMicroCT) to perform a three-dimensional (3D) analysis of viviparity associated morphological adaptations of tsetse female reproductive tract. These include 1) abdominal modifications facilitating the extreme abdominal distention required during blood feeding and pregnancy; 2) abdominal and uterine musculature required for parturition of developed larvae; 3) reduction of ovarian structure and capacity; 4) structural features of the spermatophore form in the female uterus to enhance semen/sperm delivery and inhibition of insemination by competing males; 5) uterine morphological features facilitating expansion and contraction before, during and after pregnancy; 6) milk gland structural optimizations facilitating nutrient incorporation and transfer into the uterus. The use of pcMicroCT provides unprecedented opportunities for examination and discovery of internal morphological features not possible with traditional microscopy techniques and new opportunities for comparative morphological analyses over time and between species.

## Introduction

Tsetse flies (genus *Glossina*) are the sole vectors of African trypanosomes, the causative agents of Sleeping Sickness in humans and Nagana animals. These are fatal diseases predominantly affecting marginalized populations in sub-Saharan Africa which cause severe health and economic impacts in affected countries^1^. Vector control methods are a primary method of control as tsetse flies have a low reproductive capacity relative to other insects ^2^. This is due to specializations in tsetse’s reproductive biology that results in the production of a small number of offspring over a long period of time. Tsetse females reproduce by obligate viviparity which is defined as having intrauterine larval development and provision of all larval nourishment for the duration of its development ^3–5^. Each gonotrophic cycle lasts 10 days, which restricts progeny per female to 8-10 during their lifespan.

Mating and transfer of seminal fluid into the uterus stimulates behavioral and physiological changes in female tsetse flies. Within 48 hours of mating, females become refractory to additional mating attempts by other males, start taking larger blood meals and begin to ovulate. This physiological response is exploited for control purposes via use of sterile insect technique ^6,7^. However, in *Glossina*, little is known regarding the biology underlying the post mating response.

How the post mating response in *Glossina* is triggered by mating associated stimuli and what happens to internal female morphology in response to these stimuli remains unknown. The reproductive tissues of tsetse flies are highly derived relative to that of other dipterans. The most dramatic of the reproductive modifications include the reduction of ovarian capacity ^9^, enhanced uterine musculature ^10^ and modification of the female accessory gland into a tubular ramified milk-producing organ ^11–13^. Descriptions of *Glossina* reproductive tissues have been made via a variety of media including detailed hand drawn illustrations of dissected tissues as well as via various light and electron based microscopy techniques^5,11,13–20^. Most of these techniques require compromises in one form or another which can limit the scope of the analysis.

Most microscopy techniques are destructive to different extents in that they require the sample be dissected and/or cut into thin sections for staining and imaging. The process of dissection, fixation and sectioning can cause mechanical disruption of tissue shape and positioning within the insect. These processes can also damage delicate features and interactions within and between tissues. Most microscopy-based imaging generates two-dimensional images. The lack of the third dimension confines the context and scope of the information contained within the image and can limit its informative capacity. While some light microscopy-based techniques such as confocal laser scanning microscopy have the capability to generate stacked three-dimensional (3D) images. Howevver, this technique is restricted by sample section thickness/opacity which limits the potential sample thickness to ∼20-50 *µ*m and is not applicable at the level of whole mounted tissues ^20^.

X-ray Micro Computed Tomography (MicroCT) provides an alternative that addresses many of the described issues ^21^. This technique uses an X-ray beam to virtually section through a sample across 180° of rotation resulting in an image stack representing the entire volume of the specimen. The high energy of X-rays allows biological samples of any thickness to be visualized and does not result in structural damage to the sample, which allows for repeated scans or further use of the sample in alternative visualization techniques. The primary challenge of using this technique is generation of contrast in soft tissues associated with biological samples so that features of interest can be visualized in detail. With traditional microCT this requires sample fixation followed by staining with a metallic agent such as iodine ^22^. The iodine functions to absorb energy from the X-ray beam to boost contrast and define soft tissue features by absorbance.

Alternatively, the use of a system where the energy of the X-ray beam can be adjusted to provide optimum contrast is ideal. These conditions are available at synchrotron-based facilities which allow for tuning of the X-ray beam energy. In addition, synchrotron based microCT allows for imaging via differential phase contrast rather than traditional absorbance. Phase contrast microCT (pcMicroCT) facilitates visualization of phase changes in the X-ray beam resulting from its passage through materials with different refractive indices. This allows for high contrast 3D visualizations of sample composition with or without metallic staining ^23^.

Use of microCT in the analyses of female reproductive tissues in *Drosophila melanogaster* revealed that mating induces morphological changes in female reproductive tissues over the course of the post mating transition ^24^. These changes include looping/unlooping of the uterus and oviduct, repositioning of reproductive tract within the abdomen and mating induced tissue damage resulting from the male reproductive organs. Its hypothesized that the tissue damage may allow male seminal secretions access to the hemocoel of the female. Post mating morphological changes are also observed in the oviduct which connects the ovaries to the uterus. Prior to mating, the lumen of the oviduct is closed and appears to be a solid cylindrical tube of muscle and epithelial cells. Following mating the oviduct undergoes a developmental program resulting in the formation and opening of the duct lumen facilitating the passage of oocytes from the ovaries into the uterus ^25^.

In this work, the abdominal tissues from whole freshly mated female tsetse flies are imaged via pcMicroCT to evaluate the capacity of this technique to perform 3D analyses of tsetse reproductive morphology and for future comparative analyses of post-mating changes. Analysis of the resulting data has provided detailed 3D visualizations of external and internal morphological features and provides new insights into the functional roles of structural features of *Glossina* reproductive organs.

## Materials and Methods

### Biological materials

Tsetse flies (*Glossina morsitans*) utilized in this analysis were obtained as pupae from the colony maintained at the Institute of Zoology at the Slovak Academy of Sciences in Bratislava, Slovakia. Flies were reared in the Tupper Hall arthropod containment level 2 insectary in the UC Davis School of Veterinary Medicine. Flies are maintained in an environmental chamber at 25°C and 75% relative humidity with 12:12 light/dark photoperiod. Flies receive defibrinated bovine blood meals via an artificial feeding system Mondays, Wednesdays and Fridays as described ^26^. Sterile defibrinated bovine blood for feeding is obtained from Hemostat Laboratories (Dixon, CA).

### Sample collection

Tsetse pupae were placed into an eclosion cage and monitored for teneral females daily. Teneral flies were anesthetized on ice and sorted into cages by sex. At five days post eclosion, individual virgin females were combined with males in mating cages. Cages were observed for mate pairing. If pairing was not observed within 10 minutes the male was removed from the cage and a new male introduced. Tsetse flies require at least one hour of pairing for completion of the transfer and formation of the spermatophore ^27^. Mate pairs lasting for less than 60 minutes were removed from the sample pool. Within an hour of mating completion, females were anesthetized on ice for sample preparation.

### Sample preparation, fixation and staining

Chilled flies were prepared for fixation by removal of the legs and wings to allow the fixative to permeate into the haemocoel. The fly was then placed into Bouins fixative solution (acetic acid 5%, formaldehyde 9% and picric acid 0.9%) and incubated overnight at room temperature. Following fixation, flies were dehydrated using a graded series of ethanol washes (10%, 30%, 50%, 70% and 95%). Each wash was performed for 1 hour at room temperature. Flies were then stained in 1% iodine in 100% ethanol for 24 hours. After staining, flies were washed 3 times in 100% EtOH for 30 mins per wash.

### Phase contrast micro computed tomography

During imaging, samples must remain in a fixed position with no movement during the scanning process to ensure proper alignment of the image stack. In preparation for imaging, fixed and stained flies were transferred into a 1.5 mL Eppendorf tube containing unscented Purell hand sanitizer (Gojo Industries, Akron OH). The specimen was gently pushed down to the bottom of the sample tube using a pipette tip. To ensure samples remained immobilized during scanning, the bottom of another 1.5 mL Eppendorf tube was cut off and pushed into the sample tube for use as a wedge. The – wedge was gently pushed down into the sample tube until the specimen was secured between the wall of the container and the wedge. Once the specimen was secured, the remaining volume of the sample tube was filled with Purell. The sample tube was modified to attach to the sample holder (chuck) in the MicroCT imaging hutch by hot gluing a wooden dowel (3 mm diameter, 20 mm length) vertically to the flat surface of the Eppendorf tube cap. The sample was imaged using a monochromatic beam of 20 kilo electron volts (keV). Images were captured through a 4x optique peter lens system and a PCO.edge CMOS detector with a 150 mm sample to scintillator distance. The resulting size of the individual sections was 2560 x 2560 px with a resolution of 72 pixels/inch and a 32-bit depth. During the imaging process, 1596 images were captured across 180° of rotation. Image stacks (TIFF format) were produced using Xi CAM ^28^ with the gridrec algorithm as implemented in TomoPy ^29^. The resulting image stack has reconstructed voxel resolution of 1.6 microns. The raw data stack is available for download from the following link – https://datadryad.org/stash/share/yH7IFkgeWtlX07qIp3Qu5DXpt7EtM_lMGv5s7wrjoBg.

### Data processing, segmentation, visualization and analysis

Data analysis and visualization was performed in the Dragonfly software package version 4.0 developed by ORS (Object Research Systems) in Montreal, Canada. Software available at http://www.theobjects.com/dragonfly. Image volumes were imported into Dragonfly and down sampled to 16-bit depth at half resolution (1280×1280) to improve computer performance during analysis and segmentation. The image volume was cropped to eliminate portions of the volume not occupied by the sample. Upon import, dataset contrast and sharpness were enhanced using the Unsharp image processing function with a kernel size of 7, standard deviation of 3 and unsharp factor of 3. Tissue segmentation and region of interest definitions were performed using a combination of algorithmic and manual methods. Images were captured and exported from within Dragonfly. Tissue volumes, surface areas and thickness mapping were calculated by measurement of segmented voxels by Dragonfly with each voxel representing 1.6 uM^3^. The associated video was generated using the Dragonfly movie maker function. The video is available for download from https://drive.google.com/file/d/1bHU2A6Fsxb_ZuJkg3gnbfAWXuQmr-TLG/view?usp=sharing.

## Results and Discussion

### Abdominal structural and cuticular adaptations for blood feeding and pregnancy

The analysis described here encompasses the analysis of the last four abdominal segments of a female tsetse fly (*Glossina morsitans*) immediately after copulation *(Figure 1)*. This region contains the entirety of the reproductive tract as well as tissues from other organ systems such as the digestive tract, abdominal musculature, respiratory system and the fat body (nutrient storage/metabolism). The presence of these other systems within the scan provides spatial context of how these tissues interact within other abdominal tissues. It also provides clues to the structural adaptations required to accommodate the massive changes in abdominal volume required during blood feeding and pregnancy. As in most blood feeding insects, tsetse flies ingest large volumes of blood relative to their body size. After mating, females on average take blood meals weighing between 50-60 mg ^30^. In addition, pregnant females accommodate fully developed 3^rd^ instar larvae equivalent in mass to themselves. Newly eclosed female flies weigh ∼18 mg and just after a blood meal a pregnant female can weigh over 90 mg prior to water elimination via diuresis^30^. Thus a 5-fold change in mass occurs over the course of a pregnancy cycle. It follows that to accommodate these dynamic changes, abdominal volume can increase by up to 60%.

**Figure 1:**
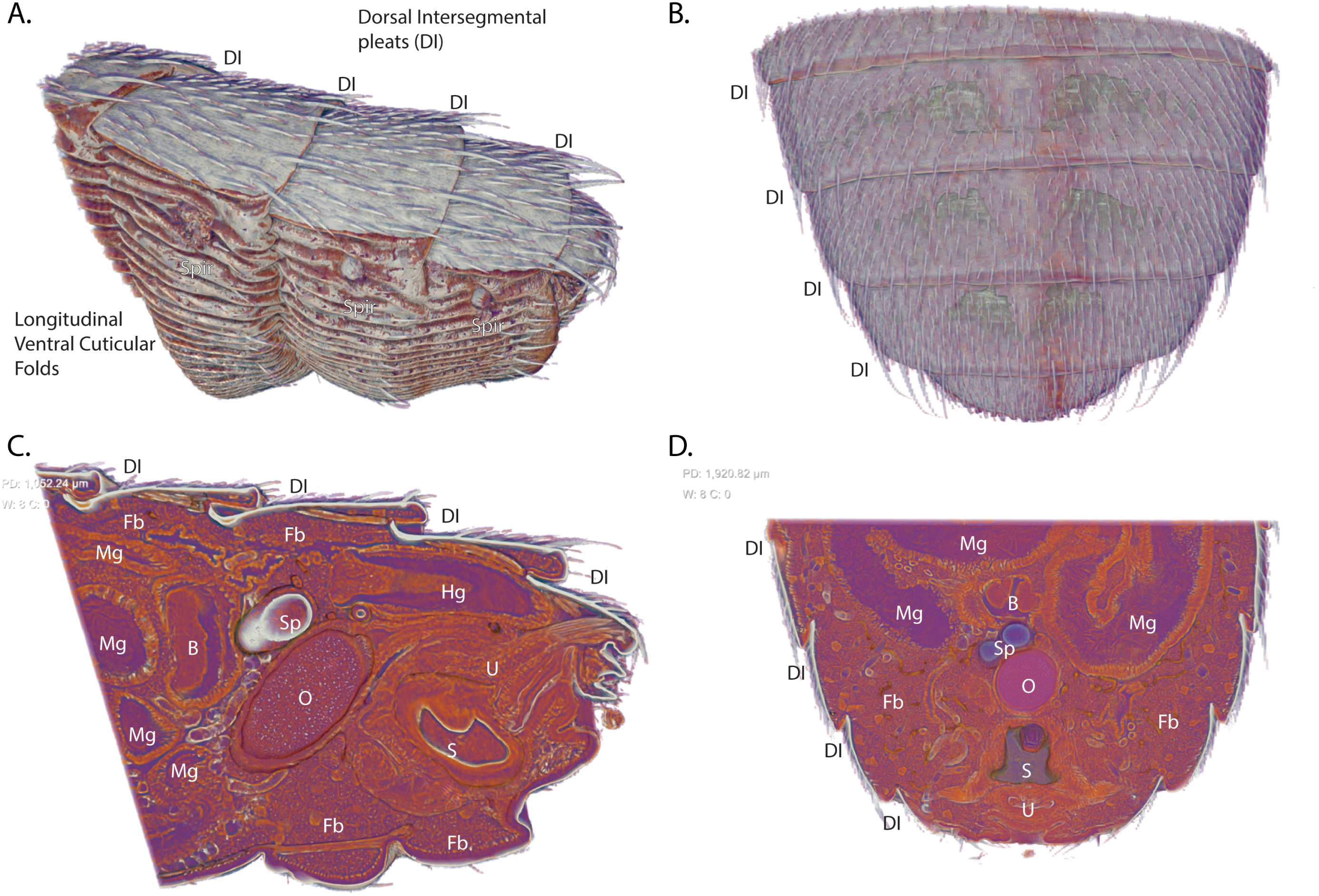
Overview of the scanned abdominal volume of the last four abdominal segments of a female tsetse fly (*Glossina morsitans*) 2 hours post mating. **A.** Side view. **B.** Top view. **C.** Sagittal Section. **D.** Coronal section through the volume. **Abbreviations:** DI – Dorsal intersegmental pleats, Fb – Fat Body, Mg – Midgut, B – Bacteriome, Sp – Spermatheca, Hg – Hindgut, O – Ovaries, U-Uterus, S – Spermatophore, Spir – Spiracle

A detailed view of exterior (Figure 1A+B) and interior (Figure 1C+D) structural features reveals some of the adaptations that facilitate abdominal expansion and elasticity. The dorsal surface of the abdominal cuticle forms pleats (dorsal intersegmental pleats) at the junction of the abdominal segments. The dorsal cuticular plates are sclerotized and covered in setae while the cuticle connecting the plates is unsclerotized and pliable. This connective cuticle folds under the proceeding segment allowing the dorsal surface of the abdomen to expand and contract along the length of the abdomen. The pleats observed on the dorsal surface are not present on the ventral surface of the abdomen. The cuticle on ventral side is soft and pliable resembling the connective cuticle associated with the dorsal plates. The ventral cuticle is marked by longitudinal folds that, along with the cuticle pliability, allow it to accordion out and expand to accommodate large changes in volume. The structure of the ventral abdominal folds bears functional similarities to the ventral pleats of rorqual whales that feed on krill and schools of small fish. These whales also undergo huge changes in volume as during feeding as they swallow volumes of water equivalent or larger than themselves ^31^.

The sagittal and coronal sections through the abdomen reveal the how tightly packed the abdominal organs are in this space (Figure 1C+D). The posterior volume of the abdomen is dominated by the ovaries, uterus and spermatheca with the hindgut traveling over the top of the reproductive tract. The anterior section of the abdomen is primarily occupied by the midgut which winds throughout the abdomen. The bacteriome region of the midgut which houses the obligate symbiont *Wigglesworthia* is clearly visible and positioned along the center line of the abdomen and anteriorly adjacent to the reproductive tract. The remaining volume is predominantly occupied by fat body cells, trachea and malphigian tubules.

### Functional aspects of Glossina abdominal musculature (Figure 2)

Internal examination of the dorsal segmental pleating reveals that the medial portion of each abdominal fold is connected to the dorsal plate anterior to it via an array of muscles (Dorsal intersegmental musculature – DIRM) (Figure 2A+B). Each muscle is angled medially from its anterior origin to its insertion onto the succeeding dorsal plate. The medial most DIRM of the final two segments have paramedian origins but converge to the midline at their insertions. Contraction of DIRM muscles looks to pull the intersegmental pleats in towards the plates anterior to them which would result in retraction of the dorsal abdominal segments. The angle at which the muscles are oriented suggests they aid in the birthing process by retraction of the dorsum to assist in dilating the vaginal canal. This musculature appears to function indirectly, relative to the rest of the uterine musculature, as it does not directly connect to the reproductive tract.

**Figure 2:**
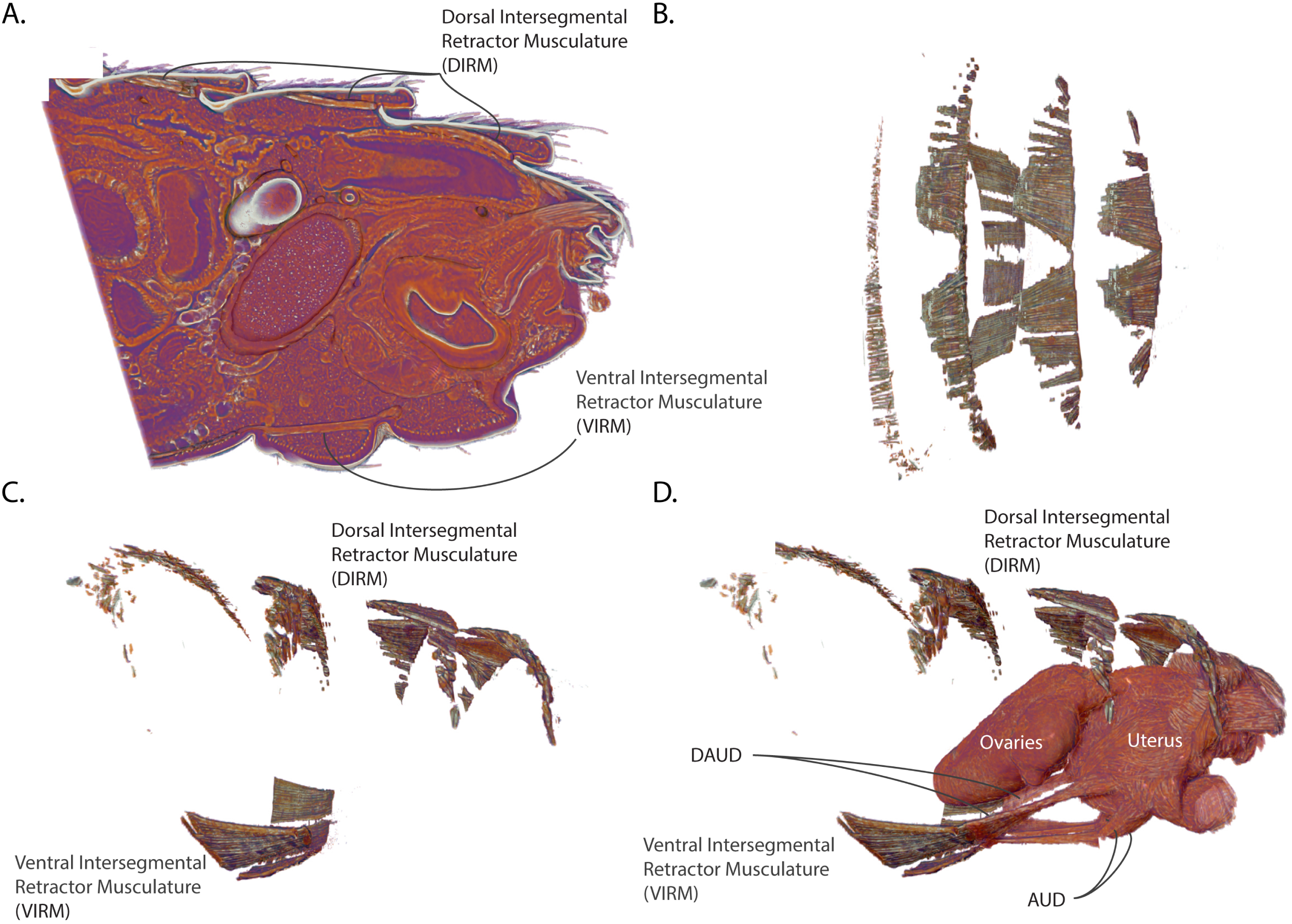
Views of the abdominal musculature and uterine connections. **A.** Sagittal section. **B.** Dorsal View. **C.** Lateral view. **D.** Lateral view of the abdominal musculature with uterus and ovaries for context. DAUD – Diagonal anterior uterine dilatory musculature, AUD – Anterior uterine dilatory musculature, DIRM – Dorsal intersegmental retractor musculature, VIRM – Ventral intersegmental retractor musculature.

On the ventral side, a wide band of muscles (Ventral intersegmental retractor musculature – VIRM) spans the 3^rd^ abdominal segment (Figure 2B+C). This band comprises two distinct pairs of longitudinal muscles: one paramedian pair and one lateral pair. Each VIRM is distally continuous with a muscle that inserts onto the anterior face of the uterus (Figure 2D). Of these uterine muscles, the lateral pair (Diagonal anterior uterine dilatory musculature – DAUD) extends diagonally from the medial VIRM toward midline of the uterus. The second set of uterine muscles (Anterior uterine dilatory muscle – AUD) extend as a pair from the ventral VIRM to insert onto the ventral side of the uterus. This set of muscles anchors the uterus to the ventral abdominal wall providing leverage to aid in dilation of the vaginal opening as well as to squeeze the anterior portion of the uterus to push the intrauterine larva towards the vaginal opening.

### The external uterine musculature is optimized for intrauterine larval development and parturition (Figure 3)

The uterus in *Glossina* is a compact structure covered in an elaborate symmetrical array of muscle pairs that anchor it to adhesion points in the final abdominal segment and the VIRM. Prior investigations into the structure of female *Glossina* reproductive physiology had documented some of these features and the nomenclature has been adapted from that publication^32^. The muscle groups were defined by manual annotation of the muscle striation patterns across the surface of the uterus. The seven uterine muscles can be divided into three groups based on their origins: (1) transverse sulcus separating third and fourth ventral segments – DAUD and AUD; (2) ventral posterior abdominal wall – PUC and AUR; (3) terminal dorsal abdominal segment – DPUD, DUD and VPU. Muscle groups 2 and 3 have extensive connections to cuticular invaginations and structures in the terminal abdominal segment.

**Figure 3:**
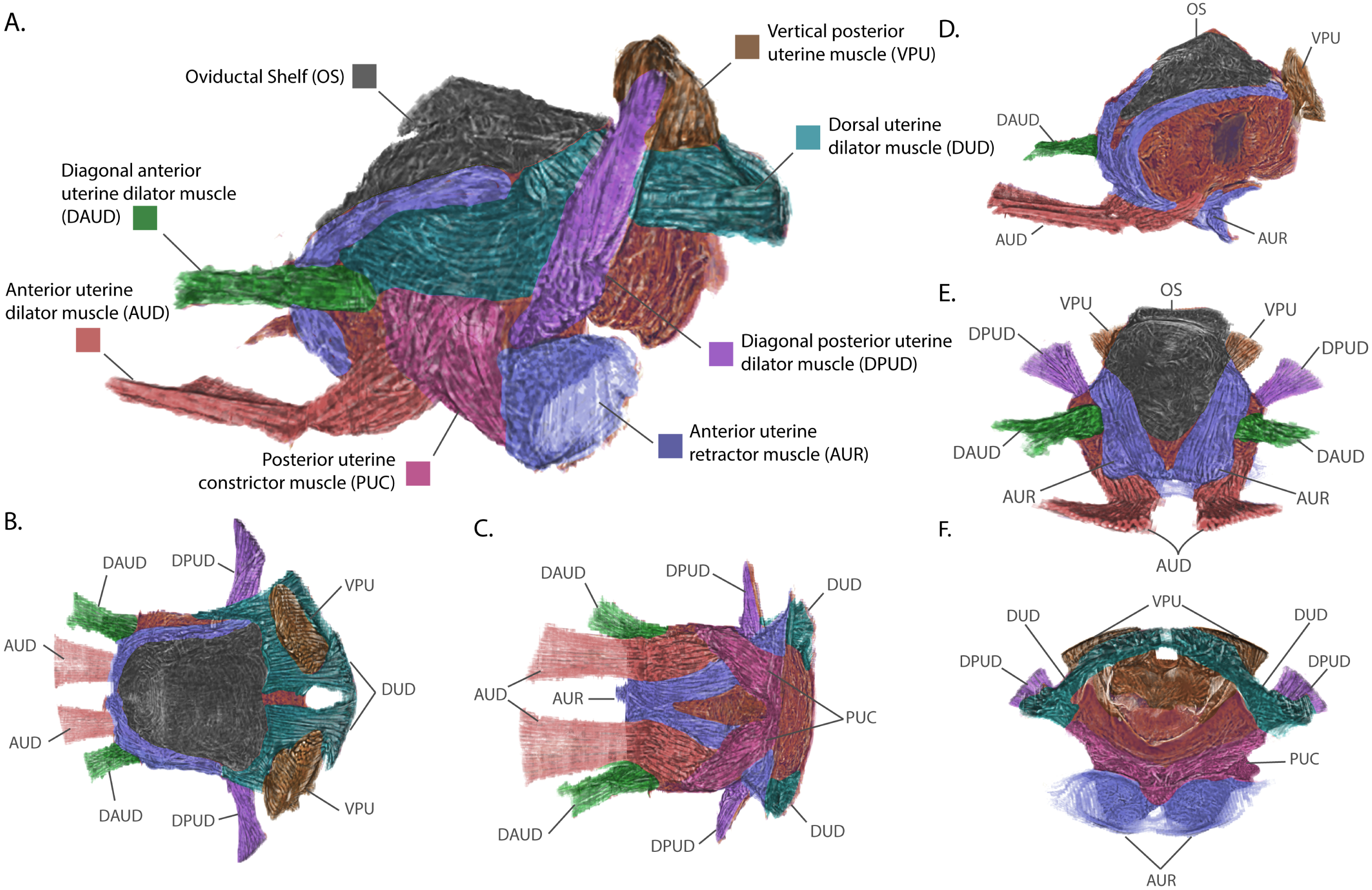
Views of the uterus and associated connective tissues with annotated musculature groups. **A.** Lateral view. **B.** Dorsal view. **C.** Ventral view. **D.** Orthogonal view with sagittal section. **E.** Anterior view. **F.** Posterior view. Muscular group colorations and acronyms are defined in the figure legend.

Among muscles originating at the ventral posterior wall, the posterior uterine dilator musculature (PUC) originates at the midline of the uterus at a point ventral to the vaginal opening. It then runs antero-laterally along the ventral and lateral sides of the uterus, passes deep to the Dorsal uterine dilator musculature (DUD) and inserts on the dorso-lateral sides of the uterus midway along its length. Due to being covered by the DUD, the insertion of the PUC is not visible in Figure 3. The PUC may aid in contraction of the uterus by providing lateral compression during parturition. The Anterior uterine retractor muscle (AUR) anchors to the surface of two bowl shaped cuticular structures that jut into the abdominal cavity from the ventral posterior abdominal wall. The AUR musculature pair cross antero-medially deep to the PUC and meet anteriorly along the ventral midline of the uterus. The AUR muscles split dorso-laterally along the anterior face of the uterus and follow the left and right sides of the ovarian shelf, finally inserting onto the postero-lateral corners of the oviductal shelf. The AUR musculature acts like straps anchoring the uterus to the posterior wall of the abdomen and can compress the anterior wall of the uterus towards the abdominal posterior. This would generate force along the antero-posterior axis to maintain uterine structure during embryogenesis/larvigenesis and aid in pushing the intrauterine larva out of the vaginal canal during parturition.

The dorsal posterior region of the uterus is anchored to the last abdominal segment by three musculature groups, the diagonal posterior uterine dilator musculature (DPUD), the dorsal uterine dilator musculature (DUD) and the vertical posterior uterine muscle (VPU). The DPUD originates on the internal dorsal surface of the last abdominal segment. It then extends ventro-medially as two bundles of muscle at a 45° angle and inserts on the posterior end of the uterus, deep to the AUR and PUC. The DUD extends anteriorly as a horizonal sheet from its origins along the posterior wall of the terminal abdominal segment. It runs parallel along the dorsal surface of the uterus inserting at the base of or underneath the ovarian shelf. The attachments made by this muscle group look to maintain close association of the posterior dorsal surface of the uterus with the last abdominal segment. The final muscle group, the VPU, originates on the inner dorsal cuticle, like the DPUD. However, the VPU origins are more medial such that the VPU extends ventrally to insert onto the dorsal surface of the vaginal opening. The muscle bundles from the VPU extend through the DUD musculature which flows around these muscles and the lateral parts of the vaginal opening prior to connection to the cuticle. The vertical positioning of the VPU suggests it also acts to secure the uterus to the last abdominal segment as well as to facilitate the opening and closing of the vaginal canal.

This analysis has revealed the basic components of the uterine musculature and additional work is in progress to compare the conformational changes these structures undergo in response to mating stimuli, ovulation and the pregnancy cycle to better understand their functional roles. Comparative analysis of *Glossina* uterine morphology with that of other viviparous flies from the Hippoboscoidea superfamily (keds and bat flies) and with that of oviparous members of the brachyceran Diptera, such as *Drosophila* and *Musca*, can inform as to the derivation of these adaptations.

### Ovarian reduction and oviduct folding reduce constraints on blood meal volume and larval size (Figure 4)

Another major adaptation to *Glossina* reproductive physiology, relative to oviparous Diptera, is the reduction in ovarian capacity. The ovaries of most oviparous Diptera contain dozens of ovarioles with some or all containing vitellogenic oocytes at the same time ^33^. In *Glossina*, each ovary contains two ovarioles with only one mature oocyte completing development per gonotrophic cycle ^8,34,35^ (Figure 4A+B). Cross section through the developing ovarian follicles shows the fully developed primary follicle with the tertiary ovarian follicle in close association inside the right ovary (Figure 4C). In the primary follicle, the oocyte has filled with yolk protein crystals and lipids and the nurse cells have emptied into the oocyte and collapsed. This oocyte has completed most of its development and is almost ready for ovulation into the uterus.

**Figure 4:**
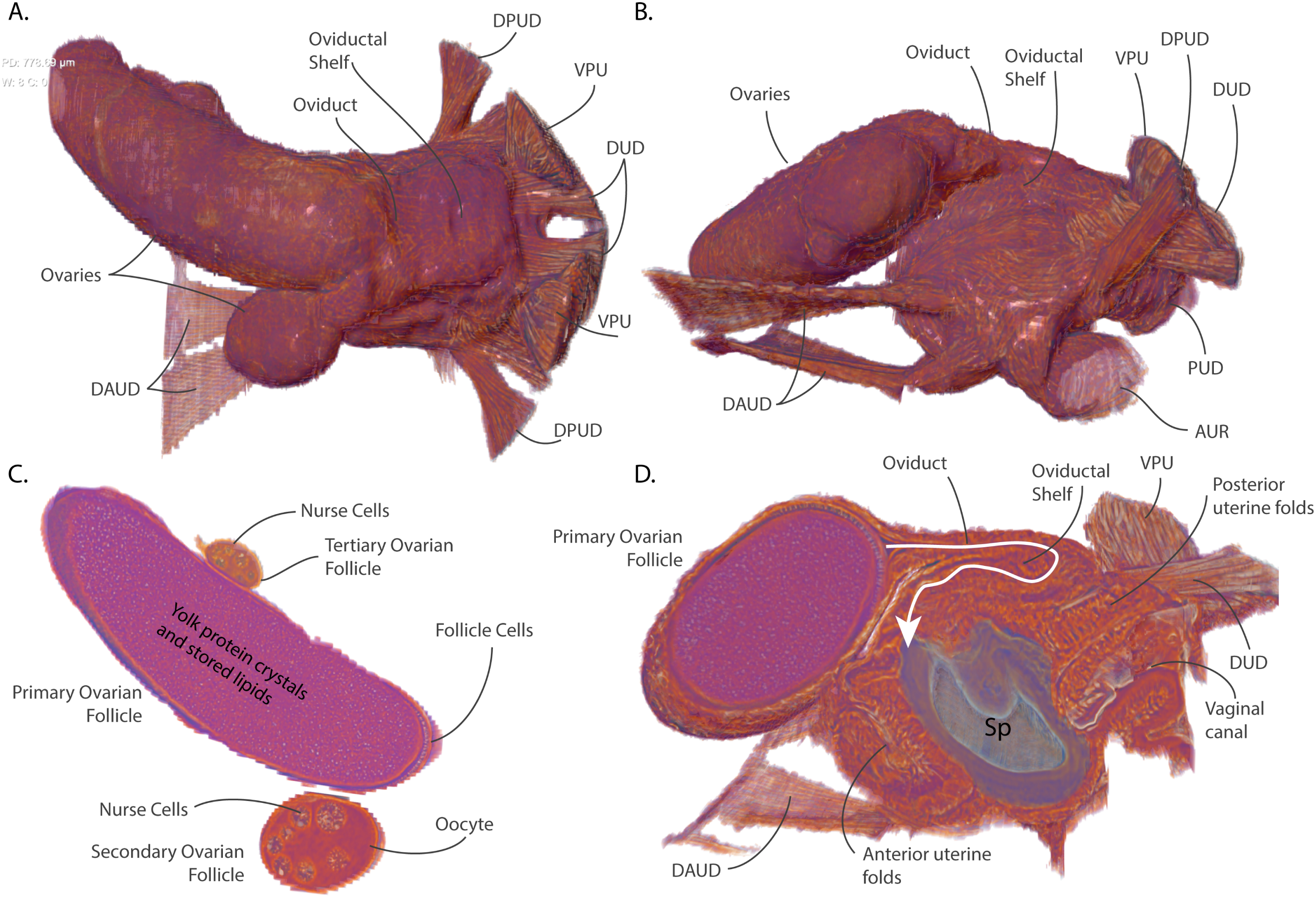
Views of the ovarian and uterine tissues highlighting ovarian structure, oocyte development and ovulation mechanics. **A.** Dorsal view. **B.** Lateral View. **C.** Coronal section of ovarian follicles. **D.** Sagittal section through the ovaries and uterus highlighting the path of the oviduct and the spermatophore inside the uterus. **Abbreviations:** DPUD – Diagonal posterior uterine dilatory musculature, VPU – Vertical posterior uterine musculature, DUD – Dorsal uterine dilator musculature, DAUD – Diagonal anterior uterine dilator musculature, AUR – Anterior uterine retractor musculature.

Tsetse flies require mating associated stimuli to undergo ovulation ^36–38^. The fly visualized in this analysis is five days post-eclosion and had just completed mating. The first ovulation usually occurs around day nine or ten post eclosion. The tertiary follicle contains nurse cells and is surrounded by follicular epithelial cells; however, it lacks a defined oocyte or any yolk deposition. The left ovary contains the secondary and quaternary ovarian follicles. The secondary follicle contains a differentiated oocyte with large nurse cells surrounded by follicular epithelial cells and a defined oocyte containing a small amount of yolk. This oocyte remains in stasis until the primary oocyte has been ovulated into the uterus. After ovulation of the primary oocyte the secondary follicle will begin rapid accumulation of yolk proteins and nutrients (vitellogenesis) in preparation for the next gonotrophic cycle. The quaternary ovarian follicle (not visible) is early in development, containing nurse cell primordia and a follicular epithelium. During the initial gonotrophic cycle, the process of oogenesis always begins with one of the follicles in the right ovary as first documented by Mellanby in 1937 ^35^. Follicular development cycles through the four ovarioles in sequence, shifting between the left and right ovaries each gonotrophic cycle.

During ovulation, the fully developed oocyte moves out of the ovary, through the oviduct and into the uterus. In *Glossina*, the oviduct and its associated tissues form a structure called the oviductal shelf which is a sleeve of tissue connecting the ovaries to the uterus (Figure 4D). The oviductal shelf folds over on itself and occupies most of the dorsal surface of the uterus. The lack of connective musculature on the dorsal anterior portion of the uterus is likely due to the spatial constraints associated with these structures. The folded conformation of this tissue results in the oviduct forming a hairpin turn prior to its opening into the uterus. The compacted structure of the oviductal shelf in addition to reduced ovarian capacity/production optimizes the space occupied by the reproductive tract within the abdominal cavity. These optimizations allow for larger blood meal volumes and increased nutrient storage capacity by the fat body between larvigenic cycles. During ovulation, the oviduct and oviductal shelf likely must expand and straighten to allow the oocyte to pass into the uterus. Whether the oviductal shelf returns to this conformation after the first ovulation and during pregnancy will need to be evaluated in scans taken later in the gonotrophic cycle.

The sagittal section of the uterus reveals that the anterior wall of the uterus also appears folded upon itself and is likely capable of expanding to accommodate the large volume required by the developing intrauterine larvae (Figure 4D). In addition, the tissue constituting the anterior and posterior uterine walls shows patterning which may represent additional folding that provides elasticity and flexibility to the uterine walls. The entire reproductive tract is optimized to conserve space while providing the capacity to expand and occupy most of the abdominal volume during intrauterine larval development when the larva is equivalent in mass to the mother.

### The spermatophore facilitates sperm storage within the spermatheca and acts as a physical barrier to insemination by competing males (Figure 5)

In *Glossina*, physical and chemical mating stimuli are required to initiate ovulation and activate other post mating changes including initiation of sexual refractoriness, accelerated oocyte development, increased host seeking/blood feeding and ingestion of increased blood meal volumes ^27,34,36,39,40^. Mating involves the transfer of a large volume of male accessory gland derived proteins and biochemicals which form a well-defined structure called a spermatophore in the uterus of the female ^41–44^. The center of this structure is hollow and contains sperm to be stored in the spermathecal organ of the female. The outer wall of this structure is composed of two layers with differing ultrastructural characteristics and has an opening at the dorsal anterior ^45^. Visualization of the spermatophore reveals that the outer walls are molded to the interior of the uterus and that it occupies a significant volume of intrauterine space (Figure 5B).

**Figure 5:**
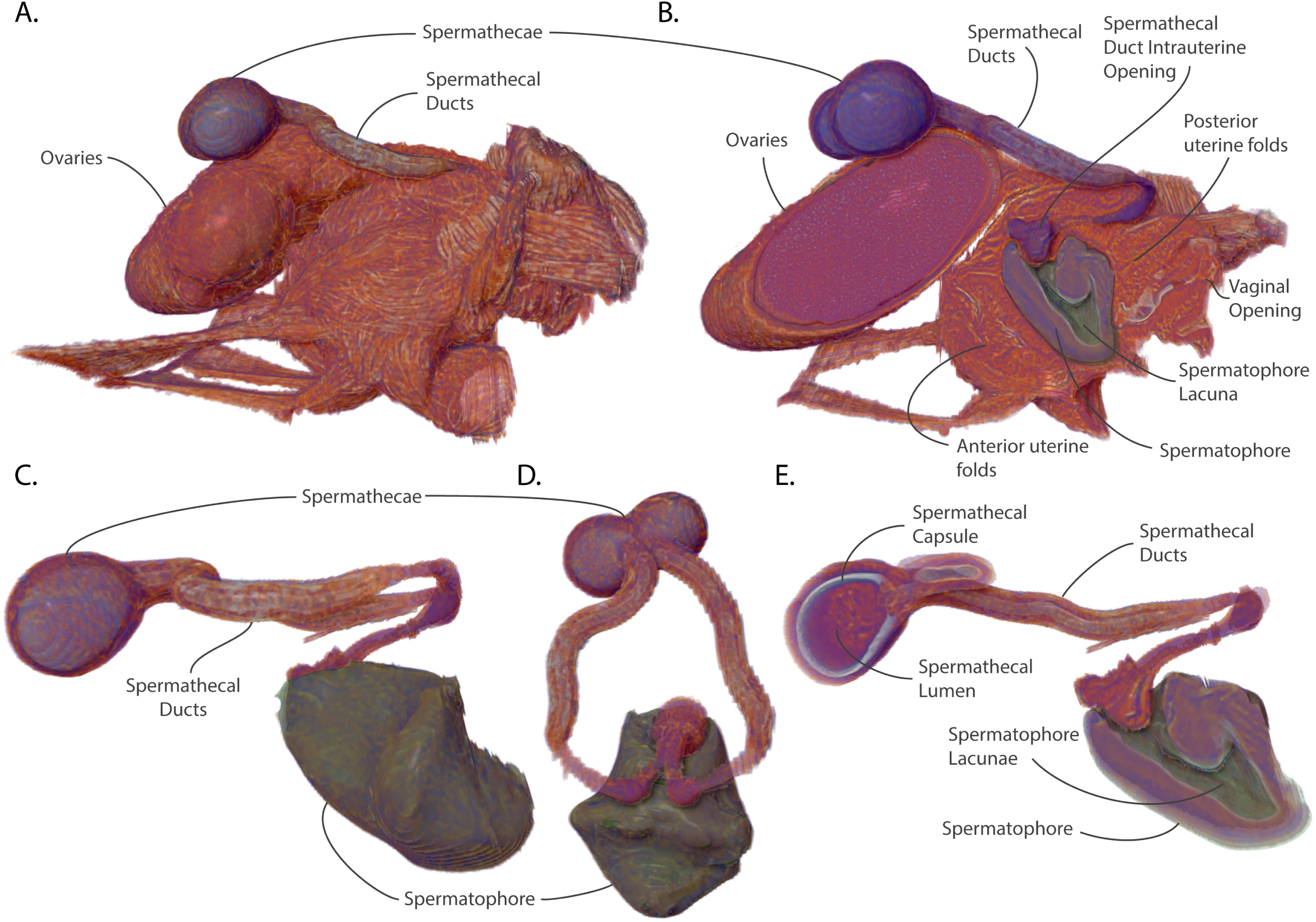
Views of the spermathecae, spermathecal ducts and spermatophore with uterus and ovaries for context. **A.** Lateral view of uterus, ovaries and spermatheca. **B.** Sagittal section of the uterus, ovaries, spermatheca and spermatophore. **C.** Lateral view of isolated spermatheca and spermatophore. **D.** Anterior view of isolated spermatheca and spermatophore. **E.** Sagittal section of spermathecae and spermatophore

The sperm storage organs, the spermathecae, and associated spermathecal ducts lie on the dorsal surface of the uterus and ovaries. The spermathecal ducts connect to the uterus as a paired tubule and share a common opening on the dorsal interior wall of the uterus (Figure 5B). From there, the paired ducts migrate posteriorly through the uterine wall and emerge from the oviductal shelf where they curve to the anterior they split into left and right tubules flanking the shelf. The tubules then rejoin dorsal to the ovaries as they open into the two spermathecal capsules (Figure 5A, C-D). The inside of the spermathecal capsule is lined with chitin and surrounding by secretory epithelial cells which secrete proteins and nutrients required to maintain the viability of stored sperm for the duration of the female’s lifespan (Figure 5E).

The lacuna of the spermatophore opens at the entrance to the spermathecal ducts in the uterus and it creates a seal against the entryway to the spermathecal ducts. This conformation guides the sperm to the opening of the ducts where they travel up to the spermathecal capsule for storage. The spermatophore also blocks access to the uterus by other males with the posterior wall inhibiting entry of seminal compounds via the vaginal canal. Females have been observed containing two spermatophores ^43^. However, the spermatophore resulting from the second mating attempt is usually stuck to the back of the first which results in a physical barrier preventing sperm from exiting the lacuna of the spermatophore. Even if the sperm were able to exit the spermatophore it is unlikely they would be able to navigate around the first spermatophore to access the spermathecal ducts.

Among the Diptera, the use of spermatophores by males is infrequent ^46^. However, in tsetse flies inter- and intrasexual competition is intense due to the low reproductive rate of females. There is evidence that there is cryptic selection by females as mating with substandard males can result in poor sperm uptake and continued receptivity to other males ^47^. Comparative analysis of male seminal protein genes between six *Glossina* species revealed them to be the most rapidly evolving genes in the genome with observed differences in gene number and sequence variability ^48^. The use of the spermatophore by males may increase the probability that a mating attempt is successful in the face of female selective pressures. The structural advantages of the spermatophore for males are that it is difficult for the female to expel, it guides the sperm to their destination and inhibits insemination by sperm from competing males. Females normally dissolve the spermatophore and expel its remnants ∼24 hours post mating and become refractory to further matings ∼48 hours post mating ^41^.

### The milk gland organ maximizes nutrient incorporation via extensive ramification for increased surface area and intimate contact with fat storage tissues (Figure 6)

The final component of the reproductive tract is the milk gland. The milk gland is a heavily modified female accessory gland which is responsible for the synthesis and secretion of a protein- and lipid-rich milk like secretion. The milk gland is a dynamic organ which undergoes cyclical changes in volume throughout the reproductive cycle in correlation with milk production activity ^45,46^. In *Glossina austeni*, it is estimated that the milk gland produces 25 mg of milk secretion per gonotrophic cycle which is equivalent to the weight of an unfed adult female ^46,47^. To accomplish this feat the gland must incorporate large amounts of stored lipids and free amino acids from dietary sources ^49,50^. The milk gland does not seem capable of protein uptake from the hemolymph and instead imports amino acids which are utilized by the massive arrays of rough endoplasmic reticulum for milk protein synthesis ^43–45^. The milk gland also harbors the extracellular form of the *Glossina* obligate symbiont *Wigglesworthia glossinidius*. The bacteria live in the lumen of the gland and are transferred to the intrauterine larva during lactation ^44–46^.

**Figure 6:**
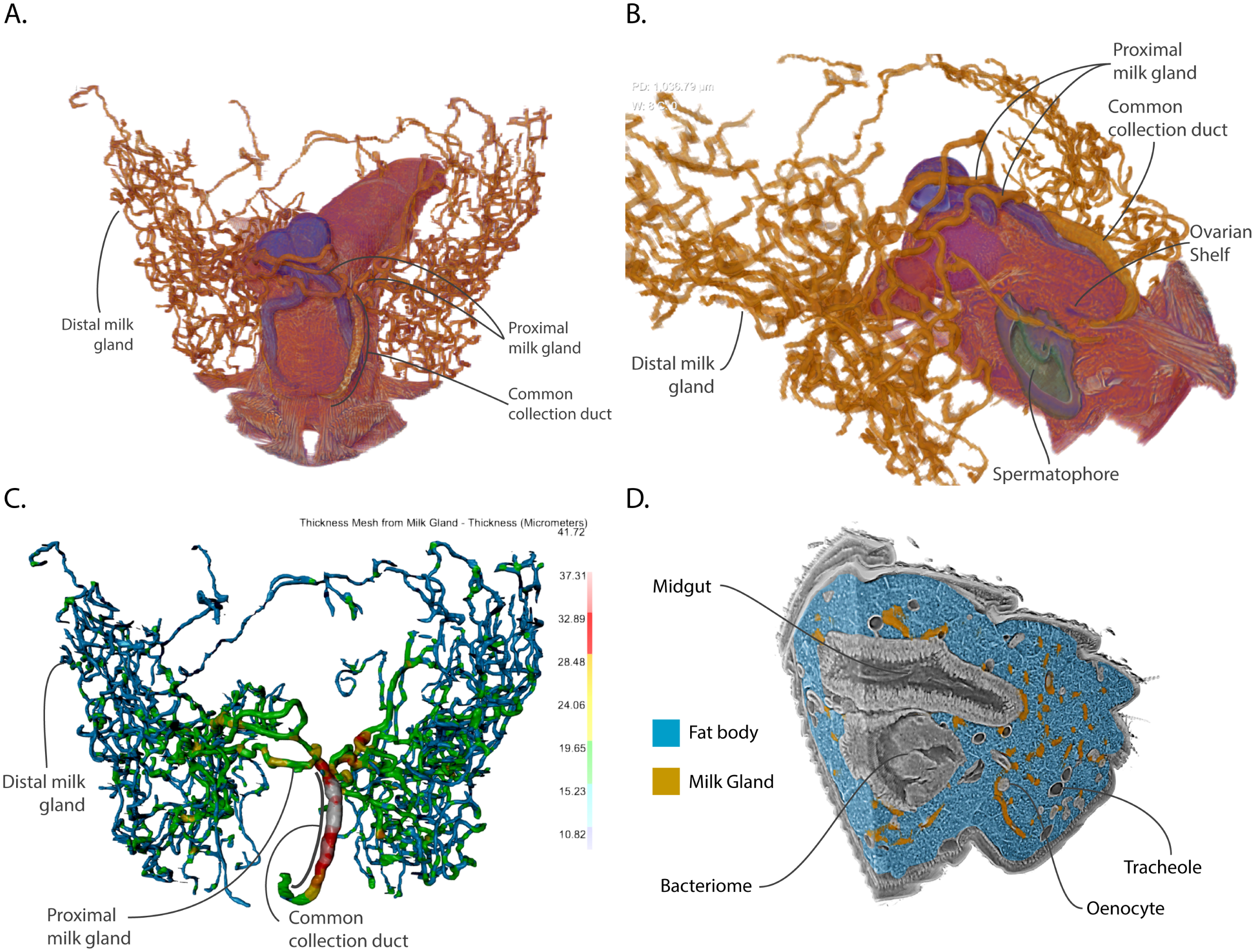
Views of milk gland morphology and structural features with other reproductive and abdominal features for context. **A.** Dorsal View. **B.** Orthologonal view with sagittal section through uterus, ovaries and spermatheca. **C.** Dorsal view of a thickness mesh of the milk gland. **D.** Orthogonal view of the abdominal volume with milk gland and fat body denoted by color.

Prior analysis of the milk gland has been primarily by 2D microscopic analysis with hand drawn interpretations of the 3D structure of this complex organ. The analysis of the abdominal volume from the pcMicroCT scan has provided a detailed virtual representation of the milk gland which clearly demonstrates the intricate nature of its branching structure (Figure 6A). The gland consists of three regions, the distal milk gland, the proximal milk gland and the common collection duct. The milk gland is connected to the dorsal surface of the uterus via the common collection duct. This duct connects to the uterus postero-ventrally to the oviductal shelf and just posterior to the opening of the spermathecal duct (Figure 6B). The duct extends through the musculature of the uterine wall and travels along the right side of the oviductal shelf where it crosses dorsally over the right spermathecal duct. The common collection duct is formed of two cuticle lined tubes lacking in secretory cells that are bundled together by a spiraling musculature, which likely regulates the flow of the milk secretions ^47–49^. At the point where the ovaries begin, the common collection duct branches into four thick tubules. Two of these cross dorsally over of the spermathecal ducts and spermathecae to expand into the left side of the abdomen while the other two proximal tubules extend to the right. The proximal milk gland lacks the musculature that wraps around the common collecting duct and contains secretory cells that contribute to milk production. The proximal milk gland rapidly transitions into the distal milk gland, which is the dominant part of the milk gland in terms of space occupied in the abdomen. The distal milk gland primarily consists of secretory cells and epithelial cells that line the interior of the gland lumen. The distal milk gland is extensively ramified and extends throughout the volume of the abdomen with most of it found within the last three segments of the abdomen. Quantitative analysis of the thickness of the gland shows that it ranges from 10-12 *µ*m at the tips of the distal milk gland to 42 *µ*m in diameter along the length of the common collecting duct (Figure 6C). The proximal gland is of intermediate diameter ranging from between 15 to 30 *µ*m with width decreasing as it transitions into the distal milk gland. The tubules of the milk gland are known to dynamically expand in diameter over the course of the pregnancy cycle ^48–50^. This female is 5 days old and early in the gonotrophic cycle. Work on visualization of the reproductive tract throughout the pregnancy cycle is ongoing and will facilitate accurate quantitative measurements of glandular changes for comparative analyses.

An inherent aspect of the tubular structure of the milk gland is that it has a very high surface area to volume ratio. Analysis of the milk gland in the context of the surrounding abdominal tissues demonstrates that it is intimately associated with fat body cells which store large volumes of lipids for milk production (Figure 6D). The large surface area of the gland facilitates rapid transfer of stored lipids during lactation. Another interesting observation is that some of the milk gland tubules are proximal to the bacteriome. The bacteriome is constituted of bacteriocyte cells that house intracellular *Wigglesworthia* and forms a horseshoe like structure that expands into the anterior midgut^51^. The mechanism by which *Wigglesworthia* invade the milk gland remains unknown and the bacteria are not found in any other tissue in the fly. Higher resolution scans of the bacteriome could provide additional information on potential structural associations between these tissues and whether *Wigglesworthia* is capable of movement between these two organs.

### Summary (Figure 7)

This analysis of the abdominal region of a newly mated tsetse fly provides a tremendous amount of information and allows the viewer to observe the spatial relationships between tissues with diverse functions that work synergistically to achieve the overarching goal of reproduction (Figure 7A-C). Analysis of the respective volume and surface area relationships reveals tissue optimizations associated with functional roles (Figure 7D). The reproductive tract (including the ovaries and uterus) comprises 7.33% of the abdominal volume but has a relatively low surface area to volume ratio revealing its spatial efficiency. In contrast, the milk gland and muscle tissues only comprise ∼1% of the abdominal volume yet have 4 and 2.5 times the surface area to volume ratio, respectively. This difference reflects the functional necessities of these tissues. The milk gland requires abundant contact with surrounding tissues to optimize nutrient transfer. The higher surface area of the muscle tissue results from the banding across the dorsal and ventral abdominal surfaces. The large surface area allows the musculature to distribute force evenly across the interior of the abdomen wall. This dataset has also been used to generate an animated video of the major features of the reproductive tract which provides the information in an informative, engaging and aesthetically pleasing manner (Supplemental video 1).

**Figure 7:**
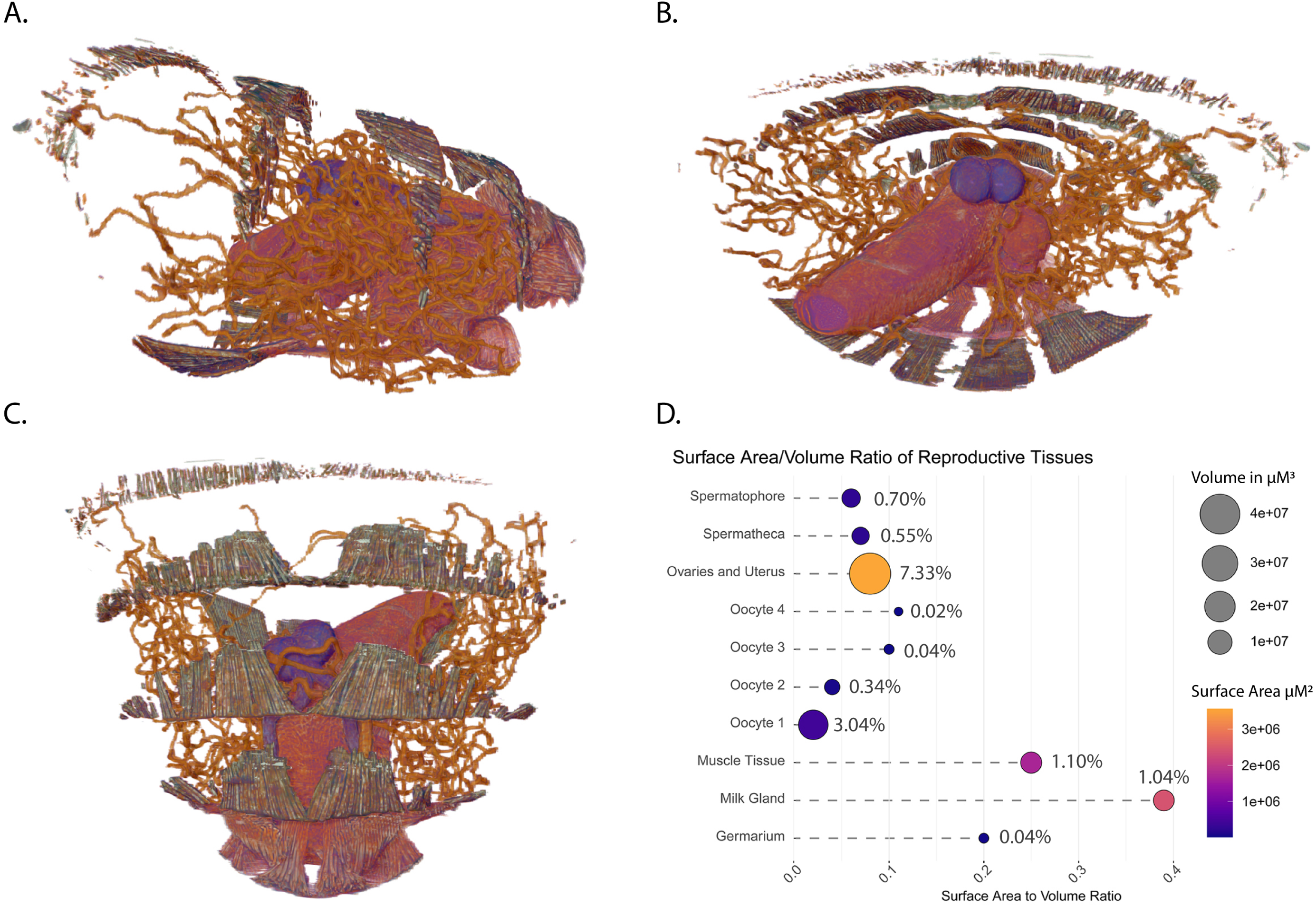
Overview of all reproductive tissues and musculature with analysis of respective volume to surface area relationships between tissues. **A.** Lateral view. **B.** Anterior view. **C.** Dorsal view. **D.** Quantification and visualization of relative volumes, surface areas, surface area to volume ratios and percentage to total abdominal volume within the scan. Circle sizes represent volume in *µ*m^3^, circle color represents surface area in *µ*m^2^, numeric annotations in percentages represent proportion of the total volume of the abdominal tissues included within the scan.

The capabilities of pcMicroCT provide tremendous opportunities for the study of internal soft tissue morphology. The non-invasive nature of this technique allows the observation of delicate relationships between internal features without the disruption caused by dissection and sectioning techniques. Once preserved, samples can be stored for years and multiple scans to be taken of the same sample at different resolutions or in different regions to highlight other morphological aspects. The ability to view external and internal structures from any angle and then to isolate and view specific features in three dimensions provides additional context, aids in viewer comprehension and facilitates deeper analysis of the data. Another tremendous benefit to this technique is that scanned volumes can be made publicly available similarly to datasets from other high throughput technologies. Users can download these volumes for independent analysis of features that were not a focus of the original analysis. Finally, the 3D nature of this data allows export of it to new 3D visualization technologies such as virtual reality applications. With the aid of a virtual reality headset, the user can explore the physiology interactively in 3D at an otherwise impossible scale. The addition of spatial cues with rich visuals leverages the natural capabilities of the human brain to rapidly interpret and remember information provided in this format ^52–54^.

Still, there are significant limitations associated with this technology. The availability of facilities capable of pcMicroCT are, for the most part, limited to those associated with a synchrotron. The scans used in this study were performed at the Lawrence Berkeley National Laboratory Advanced Light Source (ALS) on Beamline 8.3.2. The ALS is funded by the United States Department of Energy. Access to beamtime is provided to users on a grant-based system. However, new lab based MicroCT technologies are available or are in development from companies such as Zeiss and Bruker that have or will integrate phase contrast capabilities.

A powerful feature of traditional microscopy is the ability to visualize gene expression patterns or protein localization via techniques such as *in situ* and immunohistochemical staining. Protocols for performing these types of analyses via microCT are in their infancy and are complicated by accessibility and even diffusion of staining and washing reagents in a whole mounted specimen. In addition, traditional staining reagents do not absorb high energy X-rays making them invisible to this imaging technique. However, metallic staining reagents coupled to horse radish peroxidase reactive substrates show promise as a way to address these issues ^55^.

Finally, efficient analysis of the large volumes of data generated by these scans is hindered by the time required for accurate annotation. Segmentation of soft tissues with similar x-ray absorbances is difficult if not impossible for automated algorithms. Phase contrast helps with this but may still be inadequate for basic contrast-based segmentation programs. The tissue segmentations performed for this volume were done mostly by hand with aid from contrast-based analysis tools and predictive algorithms. The results often require a human eye to correctly determine boundaries between adjoining tissues. New methods based on cutting edge artificial intelligence techniques such as machine learning and deep learning are in development. The software used for this analysis Dragonfly version 4.1 for Windows implements deep learning algorithms that utilize manually segmented datasets to train the algorithm which can then be used to accurately segment raw datasets with minimal human intervention. These methods are still in early stages, but will develop rapidly as more data becomes available, computing power increases and more people develop training sets ^56^. This technique was recently used to visualize the interactions between *Cordyceps* fungus interactions with the brain and musculature of carpenter ants ^57^. The segmentations generated by this analysis will be used as training data with which to segment tissues from scans at different points post mating to identify morphological changes associated with the post-mating response. The benefits of MicroCT studies for physiological and morphological analyses are clear and it is a powerful tool that will provide a novel way to approach morphological and physiological studies of vector biology and vector parasite interactions.

## Author Roles

GA – Project conception, imaging, segmentation, data analysis, manuscript preparation; NT – Tsetse rearing, sample collection, sample preparation, manuscript editing; DYP – Sample scanning, microCT training and data analysis; LKM – Sample preparation, scanning and data preparation; XJZ – Sample preparation, scanning, manuscript editing; JA – Data analysis, segmentation, manuscript editing; PT – provision of biological materials and tsetse fly maintenance; ARM – Project conception/management, manuscript editing.

## Funding

This work was generously funded by a grant from the National Institutes of Health – NIAID #1R21AI128523-01A1 (Unraveling Intersexual Interactions in Tsetse) to GA.

Time on beamline 8.3.2 at the Lawrence Berkeley National Laboratory Advanced Light Source was provided by a GUP grant from the Department of Energy ALS-11046 – Comparative Visualization of Reproductive Tissue Ultrastructure across Fly Species.

This work was partially supported by the Slovak Research and Development Agency under contract no. APVV-15-0604 entitled “Reduction of fecundity and trypanosomiasis control of tsetse flies by the application of sterile insect techniques and molecular methods.”

This research was conducted under the IAEA CRP project “Enhancing tsetse fly refractoriness to trypanosome infection(D42015)” to ARM

## Captions

**Supplemental Video 1:** This video provides an overview of the abdominal volume detailed in this analysis and provides a visual tour of the major tissues and associated features in reproductive tract of a female *Glossina morsitans* at 2 hours post-mating. The video file is available in .avi format from the following link – https://drive.google.com/file/d/1bHU2A6Fsxb_ZuJkg3gnbfAWXuQmr-TLG/view?usp=sharing

**Supplemental Data 1:** The raw data volume is available for download as a 32-bit tiff stack from the link below. The file is compressed in a .zip format and includes 1596 sequentially numbered .tiff files – https://datadryad.org/stash/share/yH7IFkgeWtlX07qIp3Qu5DXpt7EtM_lMGv5s7wrjoBg

